# Single-dose replicating RNA vaccine induces neutralizing antibodies against SARS-CoV-2 in nonhuman primates

**DOI:** 10.1101/2020.05.28.121640

**Authors:** Jesse H. Erasmus, Amit P. Khandhar, Alexandra C. Walls, Emily A. Hemann, Megan A. O’Connor, Patience Murapa, Jacob Archer, Shanna Leventhal, Jim Fuller, Thomas Lewis, Kevin E. Draves, Samantha Randall, Kathryn A. Guerriero, Malcolm S. Duthie, Darrick Carter, Steven G. Reed, David W. Hawman, Heinz Feldmann, Michael Gale, David Veesler, Peter Berglund, Deborah Heydenburg Fuller

## Abstract

The ongoing COVID-19 pandemic, caused by infection with SARS-CoV-2, is having a dramatic and deleterious impact on health services and the global economy. Grim public health statistics highlight the need for vaccines that can rapidly confer protection after a single dose and be manufactured using components suitable for scale-up and efficient distribution. In response, we have rapidly developed repRNA-CoV2S, a stable and highly immunogenic vaccine candidate comprised of an RNA replicon formulated with a novel Lipid InOrganic Nanoparticle (LION) designed to enhance vaccine stability, delivery and immunogenicity. We show that intramuscular injection of LION/repRNA-CoV2S elicits robust anti-SARS-CoV-2 spike protein IgG antibody isotypes indicative of a Type 1 T helper response as well as potent T cell responses in mice. Importantly, a single-dose administration in nonhuman primates elicited antibody responses that potently neutralized SARS-CoV-2. These data support further development of LION/repRNA-CoV2S as a vaccine candidate for prophylactic protection from SARS-CoV-2 infection.

---

Severe acute respiratory syndrome coronavirus-2 (SARS-CoV-2) first emerged in December 2019 and within 3 months, Coronavirus Disease 2019 (COVID-19), caused by SARS-CoV-2 infection, was declared a worldwide pandemic ^1–3^. Coronaviruses are enveloped, single-strand positive-sense RNA viruses with a large genome and open reading frames for four major structural proteins: Spike (S), envelope, membrane, and nucleocapsid. The S protein mediates binding of coronaviruses to angiotensin converting enzyme 2 (ACE2) on the surface of various cell types including epithelial cells of the pulmonary alveolus ^4–6^. Protection is thought to be mediated by neutralizing antibodies against the S protein ^7,8^, as most of the experimental vaccines developed against the related SARS-CoV incorporated the S protein, or its receptor binding domain (RBD), with the goal of inducing robust, neutralizing responses ^9–11^. Indeed, previous reports have shown that human neutralizing antibodies protected mice challenged with SARS-CoV ^12–14^ and Middle East respiratory syndrome (MERS)-CoV ^15^ suggesting that protection against SARS-CoV-2 can be mediated through anti-S antibodies. Additionally, SARS vaccines that drive Type 2 T helper (Th2) responses have been associated with enhanced lung immunopathology following challenge with SARS-CoV while those with a Type 1 T helper (Th1)-biased immune response are associated with enhanced protection in the absence of immunopathology ^16,17^. Therefore, an effective COVID-19 vaccine will likely need to induce, at the very least, Th1-biased immune responses comprised of SARS-CoV-2-specific neutralizing antibodies.

Nucleic acid vaccines have emerged as ideal modalities for rapid vaccine design, requiring only the target antigen’s gene sequence and removing dependence on pathogen culture (inactivated or live attenuated vaccines) or scaled recombinant protein production. In addition, nucleic acid vaccines avoid pre-existing immunity that can dampen immunogenicity of viral vectored vaccines. Recently, clinical trials were initiated with messenger RNA (mRNA) vaccines formulated with lipid nanoparticles (LNPs) and a DNA vaccine delivered by electroporation ^18^. However, mRNA and DNA vaccines may not be able to induce protective efficacy in humans after a single immunization since, similar to inactivated and recombinant subunit protein vaccines, they typically require multiple administrations over an extended period of time to become effective ^19^. Virus-derived replicon RNA (repRNA) vaccines were first described in 1989 and have been delivered in the forms of virus-like RNA particles (VRP), *in-vitro* transcribed (IVT) RNA, and plasmid DNA ^20–23^. In repRNA the open reading frame encoding the viral RNA polymerase complex (most commonly from the *Alphavirus* genus) is intact but the structural protein genes are replaced with an antigen-encoding gene ^20,24–26^. While conventional mRNA vaccines, like that initiated in a recent clinical trial, are translated directly from the incoming RNA molecules, introduction of repRNA into cells initiates ongoing biosynthesis of antigen-encoding RNA that results in dramatically increased expression and duration that significantly enhances humoral and cellular immune responses ^27^. In addition, repRNA vaccines mimic an alphavirus infection in that viral-sensing stress factors are triggered and innate pathways are activated through Toll-like receptors and retinoic acid inducible gene (RIG)-I to produce interferons, pro-inflammatory factors and chemotaxis of antigen-presenting cells, as well as promoting antigen cross-priming ^28^. As a result, repRNA acts as its own adjuvant, eliciting more robust immune responses after a single dose, relative to conventional mRNA which typically requires multiple and 1,000-fold higher doses ^29^. An effective vaccine to stop a pandemic outbreak like COVID-19 would ideally induce protective levels of immunity rapidly and after only a single dose while simultaneously reducing the load on manufacturing at scale, due to a requirement for fewer and lower doses. Since repRNA vaccines often require only a single administration to be effective ^30^, they offer considerable potential to meet this need.

Building on experience with the attenuated Venezuelan equine encephalitis virus (VEEV) TC-83 strain ^22,30–34^, we generated repRNAs incorporating sequences from the SARS-CoV-2 Spike (S) protein, including full length S (repRNA-CoV2S) (Fig. 1A). Using immunofluorescence and western blot we demonstrated efficient expression of the v5-tagged S protein in BHK cells (Fig. 1B,C). Then, utilizing convalescent serum collected 29 days after onset of COVID-19 as an immunodetection reagent, we demonstrated endogenous expression of an S protein in BHK cells, reactive with natural SARS-CoV-2 immune sera (Fig. 1C). Next, we evaluated the ability of repRNA-CoV2S to rapidly generate antibody and T cell responses in mice when formulated with a novel Lipid InOrgainc Nanoparticle (LION) designed to enhance vaccine stability and intracellular delivery of the vaccine.

**Figure 1.**
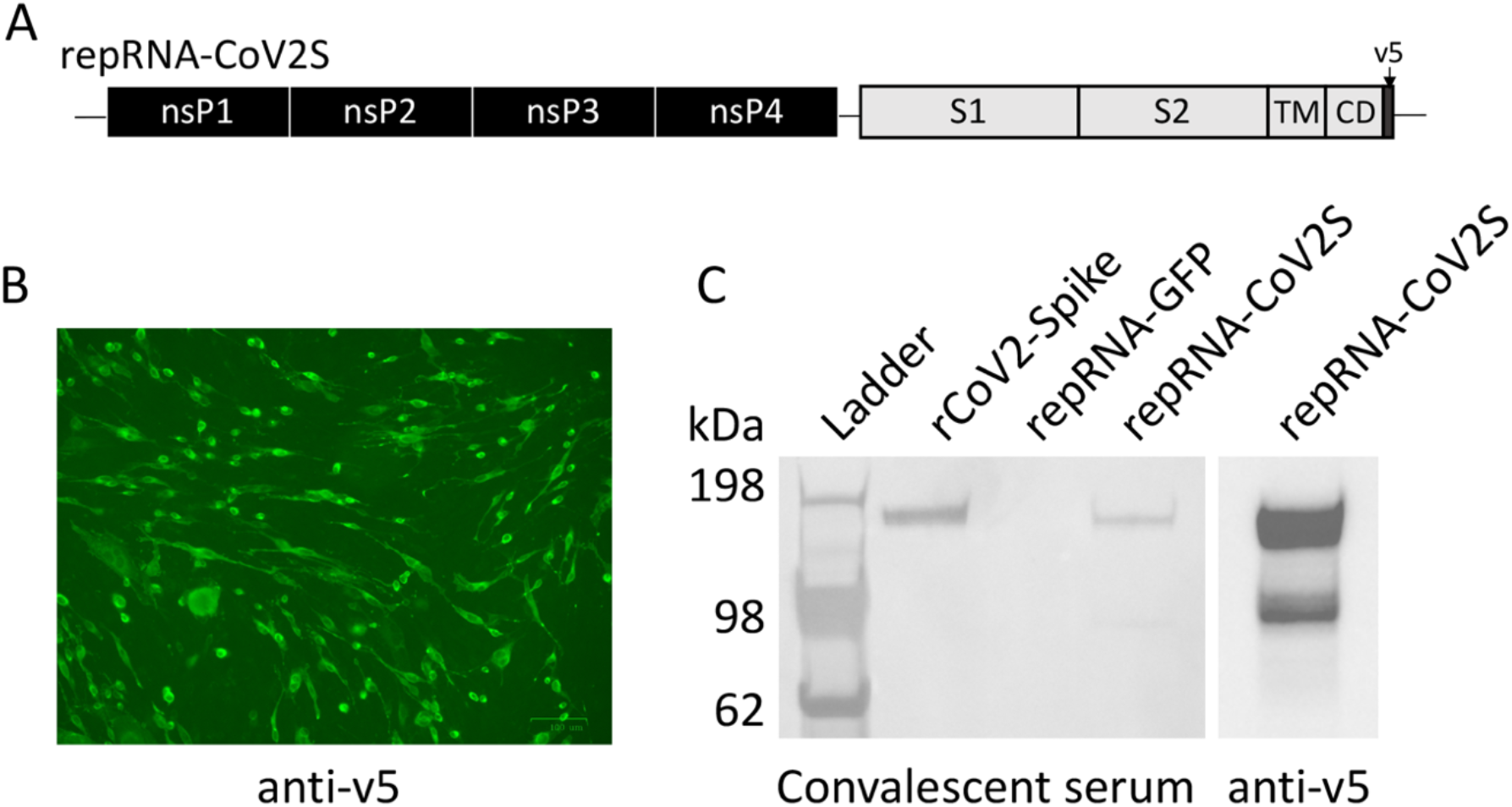
repRNA-CoV2S characterization *in vitro*. (**A**) Codon-optimized full length spike (S) open reading frame, including the S1-, S2-, transmembrane- (TM), and cytoplasmic- (CD) domains, corresponding to positions 21,536 to 25,384 in SARS-CoV-2 isolate Wuhan-Hu-1 (GenBank: MN908947.3), fused to a c-terminal v5 epitope tag, was cloned into an alphavirus replicon encoding the 4 nonstructural protein (nsP1-4) genes of Venezuelan equine encephalitis virus, strain TC-83. Following RNA transcription and capping, repRNA-COV2S, was transfected into BHK cells and 24 hours later, cells were analyzed by (**B**) anti-v5 immunofluorescence and (**C**) western blot using either convalescent human serum or anti-v5 for immunodetection. Recombinant SARS-CoV2 spike protein (rCoV2-Spike) and repRNA-GFP were used as positive and negative controls, respectively. Data in **B** and **C** are representative of 2 independent experiments.

LION is a highly stable cationic squalene emulsion with 15 nm superparamagnetic iron oxide (Fe_3_O_4_) nanoparticles (SPIO) embedded in the hydrophobic oil phase. Squalene is a known vaccine adjuvant ^35,36^ and SPIO nanoparticles have a long history of clinical use in MRI contrast and intravenous iron replacement therapy; the unique nonlinear magnetic properties of SPIOs have also been leveraged for novel use in a range of imaging, targeting and therapy applications ^37–42^. A key component of LION is the cationic lipid 1,2-dioleoyl-3-trimethylammonium propane (DOTAP), which enables electrostatic association with RNA molecules when combined by a simple 1:1 (v/v) mixing step (Fig. 2A). LION has an intensity-weighted average diameter of 52 nm (PDI = 0.2) measured by dynamic light scattering (DLS). The formulation is colloidally stable for at least 3 months when stored at 4 and 25°C (Fig. 2B). When mixed, electrostatic association between anionic repRNA and cationic DOTAP molecules on the surface of LION promotes immediate complex formation, as confirmed by increase in particle size to an intensity-weighted average diameter of 90 nm detected by DLS (Fig. 2C). Gel electrophoresis analysis of LION-formulated repRNA molecules extracted by phenol-chloroform treatment after a concentrated RNase challenge showed substantial protection from RNase-catalyzed degradation compared to unformulated repRNA (Fig. 2D). To evaluate short-term stability of the vaccine, we evaluated repRNA integrity and complex stability on 1, 4 and 7 days after mixing. LION maintained full integrity of the repRNA molecules (Fig. 2E) and complex size (Fig. 2F) at all time points.

**Figure 2.**
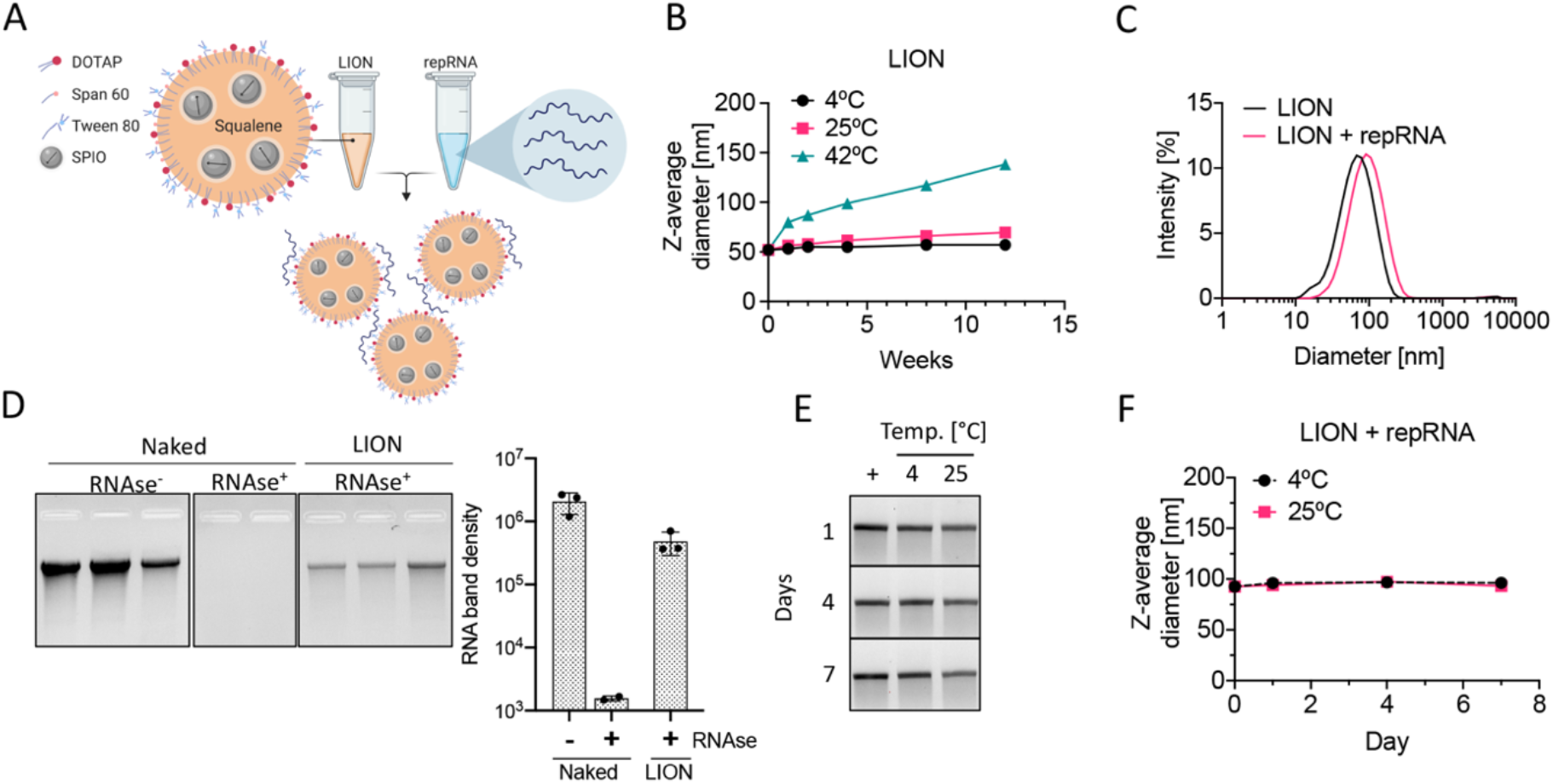
Lipid InOrganic Nanoparticle (LION) formulation of repRNA. (**A**) Graphical representation of LION and formation of vaccine complex after mixing with repRNA. (**B**) Time evolution of LION particle size, measured by dynamic light scattering (DLS), after storage at 4°C, 25°C and 42°C. (**C**) After mixing LION and repRNA, complex formation is confirmed by a shift in size distribution. (**D**) Gel electrophoresis analysis of triplicate preparations of repRNA extracted from LION after a concentrated RNase challenge shows substantial protection relative to a triplicate preparation of a dose-matched naked RNA following RNAse challenge. The formulated vaccine is stable for at least a week after mixing and storage at 4°C and 25°C as determined by (**E**) gel electrophoresis of repRNA extracted by phenol-chloroform treatment and (**F**) particle size of the complex. Data in **B**, **E**, and **F** are from a single experiment while data in **C** and **D** are representative of 3 independent experiments. Data in **B, D**, and **F** are shown as mean ± s.d. of 3 technical replicates.

A single intramuscular immunization of C57BL/6 mice with 10 or 1 μg of LION/repRNA-CoV2S induced 100% seroconversion by 14 days post-immunization and robust anti-S IgG levels with mean binding titers of 200 and 109 μg/ml, respectively, and partial seroconversion (2 out of 5) at a 0.1 μg dose (Fig. 3A). Both the 10 and 1 μg prime-only doses induced neutralizing antibodies with mean 50% inhibitory concentrations (IC50) of 1:643 and 1:226, respectively, as measured by pseudovirus neutralization assay (SARS-CoV-2 Wuhan-Hu-1 pseudotype). While all doses induced Th1-biased immune responses indicated by significantly higher IgG2c responses when compared to IgG1 (Fig. 3C), there was a trend toward higher doses inducing even more Th1-biased responses as indicated by higher IgG2c:IgG1 ratios (Fig. 3D). Given the potential role for T cells to contribute to protection, as seen with SARS and MERS ^43–45^, especially in the presence of waning antibody and memory B cell responses, we also evaluated T cell responses to LION/repRNA-CoV2S in mice. On day 28 this same cohort of mice received a second immunization and 12 days later, spleens and lungs were harvested and stimulated with an overlapping 15-mer peptide library of the S protein, and IFN-γ responses were measured by enzyme-linked immune absorbent spot (ELISpot) assay. Mice receiving a 10, 1, and 0.1 μg prime/boost exhibited robust splenic T cell responses with mean IFN-γ spots/10^6^ cells of 1698, 650, and 801, respectively (Fig. 3E). Robust T cell responses were also detected in the lung and were similar between groups with mean IFN-γ spots/10^6^ cells of 756, 784, and 777, respectively (Fig. 3F). Interestingly, analysis of the specificity of peptide response showed a biased response towards the S1 domain of the S protein in the spleen (Sup. Fig. 1A) whereas responses in the lung were more broadly distributed between the S1 and S2 domains of the S protein (Sup. Fig. 1B).

**Figure 3.**
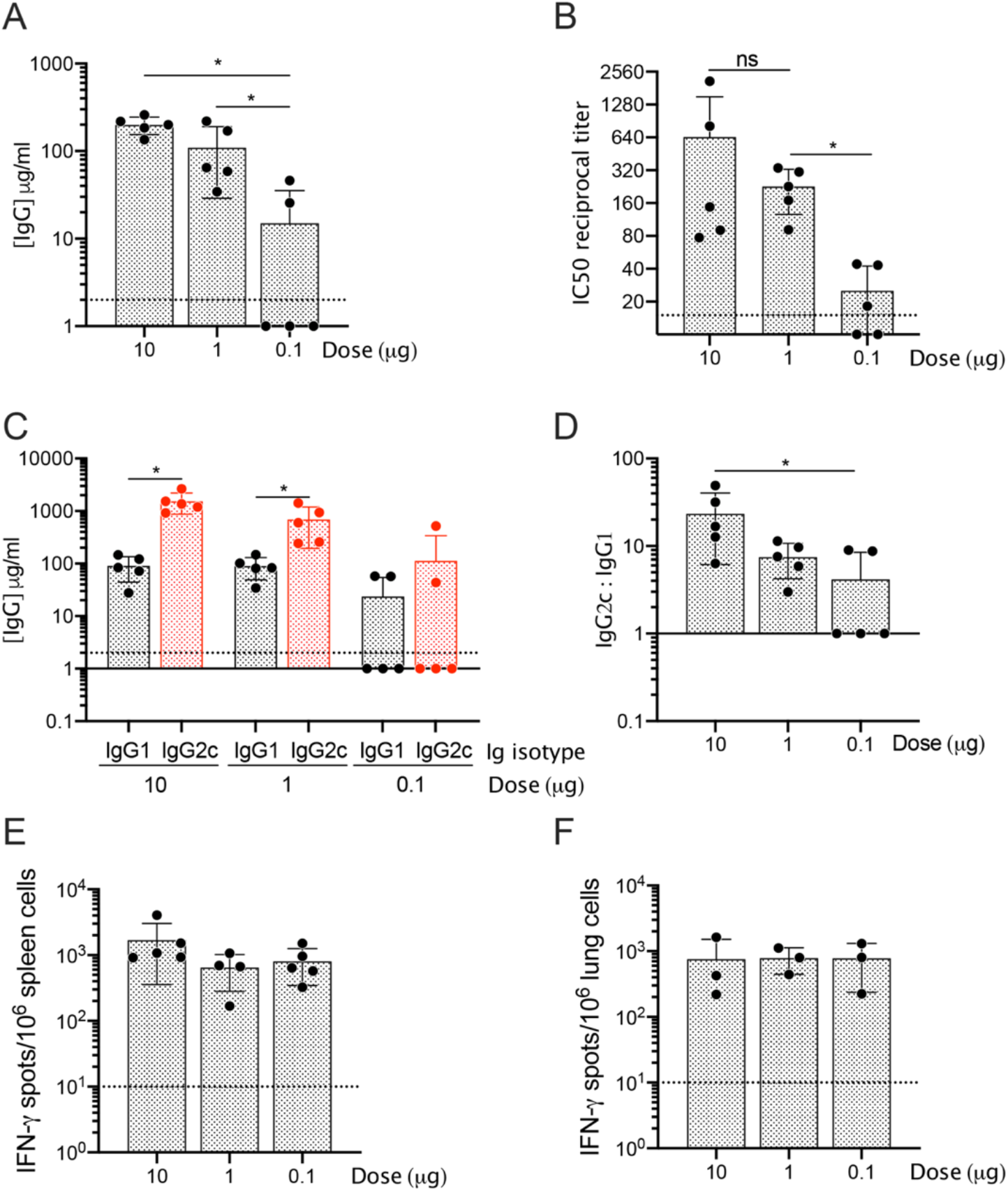
LION/repRNA-CoV2S induces Th1-biased and neutralizing antibodies in C57BL/6 mice. Six to eight-week old C57BL/6 mice (n=5/group) received 10, 1, or 0.1 μg LION/repRNA-CoV2S via the intramuscular route. Fourteen days after prime immunization, serum was harvested and (**A**) anti-S IgG concentrations were determined by enzyme linked immunosorbent assay (ELISA), (**B**) 50% inhibitory concentrations (IC50) determined by pseudovirus (SARS-CoV-2 Wuhan-Hu-1 pseudotype) neutralization assays, and (**C**) anti-S IgG1 and IgG2c concentrations and (**D**) ratios determined by ELISA. On day 28, mice received a booster immunization and 12 days later, (**E**) spleens and (**F**) lungs were harvested and IFN-γ responses were measured by enzyme-linked immune absorbent spot (ELISpot) following 18-hour stimulation with 10 peptide pools encompassing the S protein and consisting of 15-mers overlapping by 11 amino acids (see Sup. Fig. 1). Data in **A**, **C**, and **D** are representative of 3 independent experiments while data in **B**, **E**, and **F** are from a single experiment. All data are represented as individual values as well as mean ± s.d. *p<0.05 as determined by one-way ANOVA with Tukey’s multiple comparison test.

The elderly are among the most vulnerable to COVID-19 but immune senescence in this population poses a barrier to effective vaccination. To evaluate the effect of immune senescence on immunogenicity, we next administered 10 or 1 μg of LION/repRNA-CoV2S in 2-, 8-, and 17-month old BALB/C mice and measured anti-S IgG concentrations at 14 days after a single immunization. Significantly lower antibody titers were observed in the 17-month old mice at both doses (Fig. 4A), when compared to 2- and 8-month old mice, suggesting that higher doses and/or additional booster doses may be required in the most immune senescent populations to induce sufficient immunity. No differences were observed between the 2- and 8-month old mice. Interestingly, although BALB/C mice tend to develop a more Th2 immune-biased response following vaccination ^46^, LION/repRNA-CoV2S induced ratios of IgG2a:IgG1 greater than 1 (Fig. 4B, C) in all age groups of BALB/C mice, indicating a Th1-biased immune response. Given that severe, life-threatening COVID-19 appears to be more common among elderly individuals, irrespective of type of T helper response, and that severe SARS is associated with skewing toward Th2 antibody profiles with an inadequate Th1 response ^16,17,43^, the ability of LION/repRNA-CoV2S to induce strong and Th1-biased responses in 8- and 2-month old mice, even in the Th2-biased BALB/c strain, is a promising finding regarding the potential safety and immunogenicity of this vaccine.

**Figure 4.**
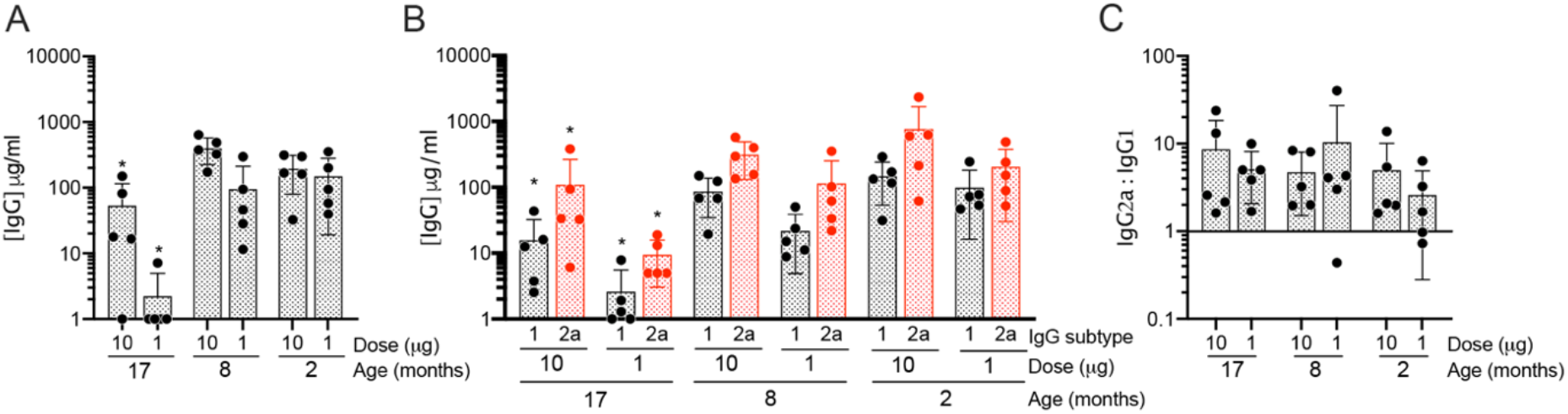
LION/repRNA-CoV2S induces Th1-biased antibodies in aged BALB/C mice. Two-, eight-, or seventeen-month old BALB/C mice (n-5/group) received 10 or 1 μg LION/repRNA-CoV2S via the intramuscular route. Fourteen days after prime immunization, serum was harvested and (**A**) anti-S IgG or (**B**) IgG1 and IgG2a concentrations and (**C**) ratios were determined by enzyme-linked immunosorbent assay (ELISA). Data in 17-, 8-, and 2-month old BALB/Cs are from a single experiment and data for the 2-month old BALB/Cs were replicated in a second experiment. All data are represented as individual values as well as mean ± s.d. *p<0.05 as determined by one-way ANOVA with Tukey’s multiple comparison test between the 17-month old animals and either the 8- or 2-month old animals.

Having achieved robust immunogenicity with LION/repRNA-CoV2S in mice, we then immunized pigtail macaques (*Macaca nemestrina*) to determine if the vaccine was capable of inducing strong immune responses in a nonhuman primate model that more closely resembles humans in the immune response to vaccination. Three macaques received LION/repRNA-CoV2S at a single intramuscular 250 μg dose at week 0 and two macaques received a 50 μg prime at week 0 and a boost at week 4. (Fig. 5A). Blood was collected 10, 14, 28, and 42 days post vaccination to monitor vaccine safety and immunogenicity. There were no observed reactions at the vaccine injection site nor adverse reactions in the animals up to 42 days post-prime vaccination. Additionally, there were no abnormalities in weight or temperature in the animals (Sup. Fig. 2A-B), and serum chemistries revealed no abnormal findings, except for mild azotemia (mildly elevated blood urea nitrogen and creatinine) in 1 animal at 14 days post vaccination (Sup. Fig. 2C). All CBC counts were unremarkable (Sup. Fig. 2D).

**Figure 5.**
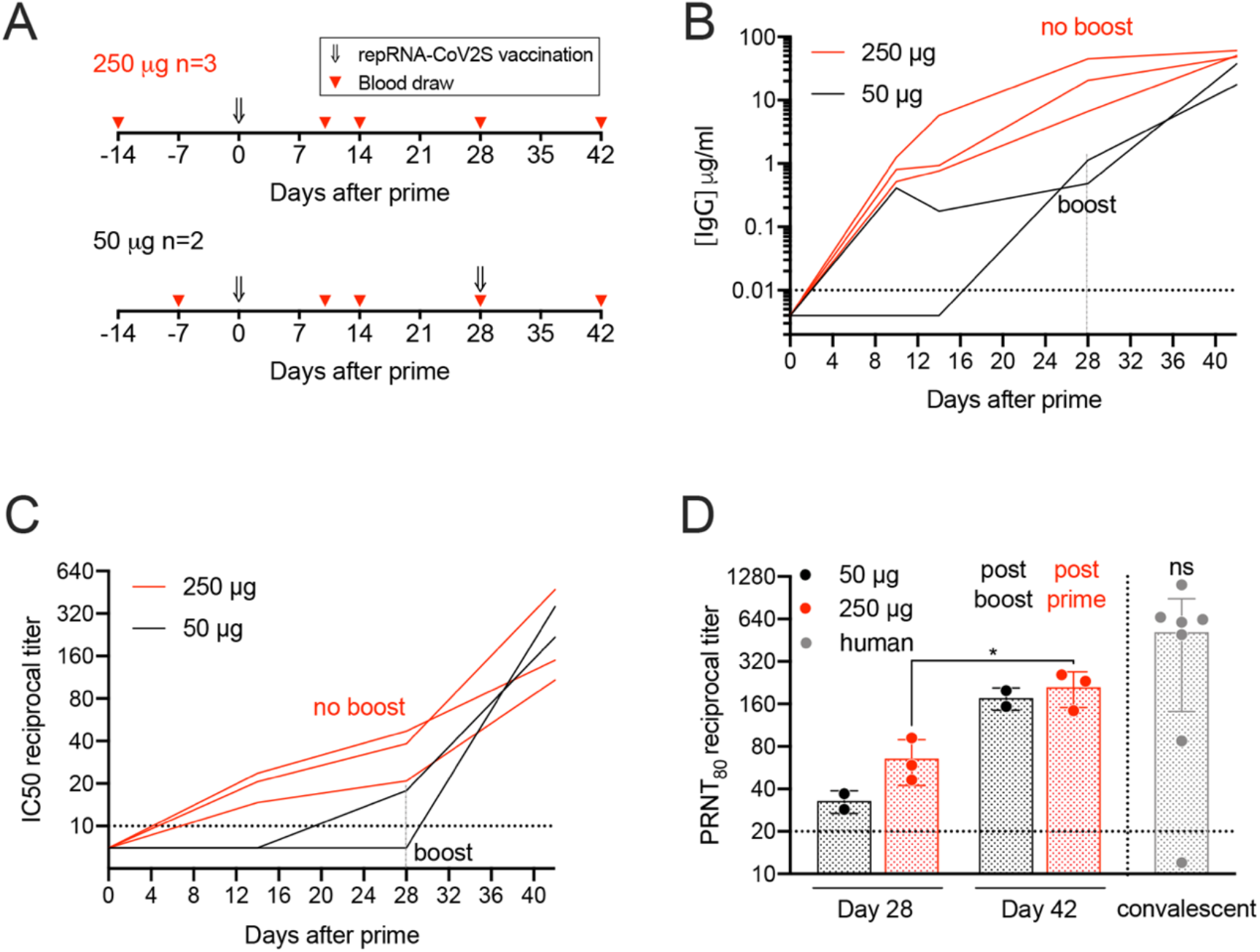
Single dose of LION/repRNA-CoV2S induces neutralizing antibody responses in pigtailed macaques. (**A**) Pigtail macaques were vaccinated with 250 μg (n=3) or with 50 μg (n=2) repRNA-CoV2-S via the intramuscular route and blood collected on days 10, 14, 28, and 42; the 50 μg group received a boost vaccination on day 28 and blood collected 14 days later. (**B**) Using pre-immunization blood draws to establish a baseline, serum anti-S IgG enzyme linked immunosorbent assays (ELISAs) were performed on the post-immunization samples as well as (**C**) pseudovirus (SARS-CoV-2 Wuhan-Hu-1 pseudotype) neutralization assays to determine mean 50% inhibitory concentrations (IC50) of each sample. Additionally, (**D**) 80% plaque-reduction neutralizing antibody titers (PRNT_80_) against SARS-CoV2/WA/2020 isolate were measured at days 28 and 42 alongside sera from 7 convalescent human samples collected from confirmed COVID-19 patients (see Sup. Table 1). The experiment was performed once. Each line in **B** and **C** are representative of each individual animal. Data in **D** are reported as individual values as well as mean ± s.d. *p<0.05 as determined by students t-test comparing 250 μg groups at days 14 and 28. There was no significant difference (ns) between mean PRNT_80_ titers in all 5 animals at day 42 and titers in sera from 7 convalescent humans, as measured by Mann-Whitney U test.

ELISA analyses (Sup. Fig. 3) of sera collected 10, 14, 28, and 42 days after prime immunization showed that all three macaques immunized with the single 250 μg dose seroconverted as early as day 10, with anti-S IgG concentrations continuing to increase in these 3 animals to 48, 51, and 61 μg/ml by day 42 (Fig. 5B). Both macaques receiving 50 μg repRNA-CoV2S seroconverted after a single dose but developed significantly lower antibody responses with anti-S IgG concentrations of 1 and 0.5 μg/ml by day 28, compared to 7, 20, and 45 μg/ml in the 250 μg group at this same time point (Fig. 5B). However, 14 days after a booster immunization, the 50 μg group developed similar levels of anti-S IgG concentrations (18 and 37 μg/ml) as the 250 μg prime-only group at this time point (48, 51, and 61 μg/ml) (Fig. 5A). Additionally, sera from the three macaques immunized with just the single 250 μg dose neutralized pseudovirus (SARS-CoV-2 Wuhan-Hu-1 pseudotype) transduction of cells *in vitro* with reciprocal IC50 titers of 1:38, 1:20 and 1:47 by day 28 with levels increasing to 1:472, 1:108, and 1:149 by day 42, whereas the 50 μg group achieved similar robust IC50 titers only after the booster immunization reaching pseudovirus IC50 titers of 1:218 and 1:358 by day 42 (Fig. 5C and Sup. Fig. 4). Sera collected 28- and 42-days post vaccination were further analyzed for neutralization of wild type SARS-CoV-2/WA/2020 by 80% plaque reduction neutralization test (PRNT_80_) and compared to neutralizing titers in sera from convalescent humans collected 15-64 days following natural infection (Sup. Fig. 4 and Sup. Table 1). A single immunization with 50 and 250 μg of LION/repRNA-CoV2S induced mean PRNT_80_ titers of 1:32 and 1:66 by day 28, respectively. By Day 42, mean PRNT_80_ titers significantly increased to 1:176 after a booster immunization in the 50 μg group and to 1:211 in the prime-only 250 μg group, (Fig. 5D and Sup. Fig. 4). Importantly, all 5 macaques developed PRNT_80_ titers within the same range as titers measured in the seven convalescent humans (<1:20 to 1:1280, collected 15 to 64 days post onset) and there was no significant difference in mean neutralizing titers between all 5 vaccinated macaques (1:197) and convalescent humans (1:518) (P=0.27, Fig. 5D, Sup. Fig. 4, and Sup. Table 1). Recently, serum neutralizing titers, measured as the IC50 titer that neutralized SARS-CoV-2 by 50% tissue culture infectious dose (TCID_50_), were reported in rhesus macaques that were either re-infected ^47^ or challenged after vaccination with an inactivated SARS-CoV-2 vaccine ^48^. In the former report, IC50 titers as low as 1:8 were associated with protection from re-infection while in the latter, IC50 titers as low as 1:50 were associated with reduced viral load and protection from lung pathology. These data suggest that a 250 μg prime-only or a 50 μg prime/boost immunization with the LION/repRNA-CoV2 vaccine may be able to induce levels of neutralizing antibodies sufficient to protect nonhuman primates from infection and disease. Studies are now underway to evaluate protective efficacy.

RepRNA vaccines against a variety of infectious diseases and cancers have been shown to be safe and potent in clinical trials ^49–52^, and the cell-free and potentially highly scalable manufacturing process of repRNA when used with effective synthetic formulations, such as LION, present further benefits over mRNA. The two-vial approach provides a significant manufacturing and distribution advantage over LNP formulations that encapsulate RNA, as the vaccine can be stockpiled and combined onsite as needed. Additionally, we demonstrated that LION/repRNA-CoV-2 induces robust S-specific T cell responses in mice. Given the relatively recent emergence of SARS-CoV-2, we can only speculate based on limited knowledge from previous reports of coronavirus infection as to how T cell responses may contribute to protection from infection and disease. Following natural infection of humans with the related SARS-CoV, neutralizing antibody and memory B cell responses in some individuals are reported to be short lived (^~^ 3 years) while memory T cells persist at least 6 years ^53^, suggesting a potential role for T cells in long term responses especially in those who lack robust memory B cell responses. Additionally, anti-S T-cell responses to the related SARS- and MERS-CoVs contribute towards viral clearance in normal as well as aged mice infected with SARS- or MERS-CoV, respectively ^43–45^.

Together, our results demonstrate a significant potential for LION/repRNA-CoV2S, which will enter clinical development under the name HDT-301, to induce rapid immune protection from SARS-CoV-2 infection. A scalable and widely-distributed vaccine capable of inducing robust immunity in both young and aged populations against SARS-CoV-2 infection in a single shot would provide immediate and effective containment of the pandemic. Critically, the vaccine induced Th1-biased antibody and T cell responses in both young and aged mice, an attribute that has been associated with improved recovery and milder disease outcomes in SARS-CoV-infected patients ^54^. Together, these results support further development of LION/repRNA-CoV2S as a vaccine candidate for protection from COVID19.

## Acknowledgements

The authors would like to thank Brieann Brown, Solomon Wangari, Joel Ahrens, Naoto Iwayama, and William Garrison for their technical assistance with the pigtail macaque study as well as Dr. Helen Chu and Sarah Bowell for donating remnant, de-identified convalescent human sera from confirmed COVID-19 patients. Additionally, the authors thank Dr. Scott Weaver at the University of Texas Medical Branch for providing the plasmid vector encoding VEEV-TC83, and the Institute for Protein Design at the University of Washington for providing recombinant SARS-CoV-2 spike protein.

This work was funded by P51OD010425 (Washington National Primate Research Center), NIH/NIAID Centers of Excellence for Influenza Research and Surveillance contract HHSN27220140006C (JHE), and HDT Biotech internal funds. Additional support from the University of Washington Center for Innate Immunity and Immune Disease, NIH/NIAID contract 75N93019C00037 (MD), NIH/NIAID contract 75N93019C00008 (APK), the NIGMS/NIH R01GM120553 (DV), NIAID/NIH HHSN272201700059C (DV), a Pew Biomedical Scholars Award (DV), an Investigators in the Pathogenesis of Infectious Disease Award from the Burroughs Wellcome Fund (DV), and the intramural research program of NIAID, NIH (DH, SL, HF). JHE is a Washington Research Foundation Postdoctoral fellow and is also supported by NIH 1F32AI136371. The content is solely the responsibility of the authors and does not necessarily represent the official views of the funders.

## Supplementary Material

### Materials and Methods

#### SARS-CoV-2 repRNA vaccine production and qualification

Codon optimized gene sequences for SARS-CoV-2 full S corresponding to positions 21,536 to 25,384 in SARS-CoV-2 isolate Wuhan-Hu-1 (GenBank: MN908947.3) fused to a c-terminal v5 epitope tag was synthesized as double stranded DNA fragments (IDT) and cloned into a plasmid vector encoding the 5’ and 3’ untranslated regions as well as the nonstructural open reading frame of Venezuelan equine encephalitis virus, strain TC-83, between PflFI and SacII sites by Gibson assembly (SGI-DNA). Clones were then sanger sequenced and prepped for RNA production as follows. Template DNA was linearized by enzymatic digestion with NotI followed by phenol chloroform treatment and ethanol precipitation. Linearized template was transcribed using MEGAscript^®^ T7 Transcription Kit (Invitrogen, Carlsbad, CA) followed by capping with NEB Vaccinia Capping System as previously described ^1^. To qualify the vaccine candidate *in vitro*, Baby Hamster Kidney (BHK) cells (ATCC) were transfected with repRNA or mock transfected using *TransIT-mRNA* transfection kit (Mirus Bio) and cells analyzed 24 hours later by immunofluorescence using a mouse anti-v5 AF488 secondary antibody (Invitrogen). Additionally, BHK cells were transfected with repRNA-CoV2S and repRNA-GFP and cell lysates were collected 24 hours later for analysis by SDS-PAGE and by western blot using recombinant SARS-CoV-2 S protein as a positive control. To detect repRNA-mediated protein expression following transfer to nitrocellulose membrane, anti-v5-HRP or convalescent human serum collected 29 days after onset of PCR-confirmed COVID-19 followed by anti-human Ig-HRP secondary antibody (Southern Biotech) was used.

#### LION formulation

To protect the RNA replicons from degradation, we partnered them with a Lipid InOrganic Nanoparticle (LION) formulation that consists of inorganic superparamagnetic iron oxide (SPIO) nanoparticles within a hydrophobic squalene core to enhance formulation stability. LIONs comprise 37.5 mg/ml squalene (Millipore Sigma), 37 mg/ml Span^®^ 60 (Millipore Sigma), 37 mg/ml Tween^®^ 80 (Fisher Chemical), 30 mg/ml DOTAP chloride (Corden Pharma), 0.2 mg/ml 15 nm oleic acid-coated iron oxide nanoparticles (Ocean Nanotech, San Diego, CA) and 10 mM sodium citrate dihydrate (Fisher Chemical). LION particles were manufactured by combining the iron oxide nanoparticles with the oil phase (Squalene, Span 60, and DOTAP) and sonicating for 30 minutes in a 65°C water bath. Separately, the aqueous phase, containing Tween 80 and sodium citrate dihydrate solution in water, was prepared with continuous stirring until all components were dissolved. The oil and aqueous phases were then mixed and emulsified using a VWR 200 homogenizer (VWR International) and the crude colloid was subsequently processed by passaging through a microfluidizer at 20,000 psi with a LM10 microfluidizer equipped with a H10Z-100 μm ceramic interaction chamber (Microfluidics) until the z-average hydrodynamic diameter – measured by dynamic light scattering (Malvern Zetasizer Nano S) – reached 50 ±5 nm with a 0.2 polydispersity index. The microfluidized LION was terminally filtered with a 200 nm pore-size polyethersulfone (PES) filter and stored at 2-8°C.

#### RNase protection

Replicon RNA was complexed with LION formulations and placed on ice for 30 min. After diluting the complex using nuclease-free water, complexes containing 1 μg of repRNA at 20 μg/mL were treated with 50 ng of RNase A (Thermo Scientific) for 30 min at room temperature, followed by an incubation with 5 μg of recombinant Proteinase K (Thermo Scientific) for 10 min at 55°C. RNA was then extracted using an equal volume of 25:24:1 phenol:chloroform:isoamyl alcohol (Invitrogen). After vortexing, samples were centrifuged at 17,000 × *g* for 15 min. The supernatant was collected and mixed 1:1 with Glyoxal load dye (Invitrogen) and heated at 50°C for 15 min. The equivalent of 200 ng of RNA was loaded and run on a denatured 150 mL 1% agarose gel in Northern Max Gly running buffer (Invitrogen) at 120 V for 45 min. Gels were imaged using a ChemiDoc MP imaging system (BioRad). The intensity of the intact RNA band was compared to phenol:chloroform:isoamyl extracted RNA from complexes that were not subjected to RNase and Proteinase K treatment.

#### Mouse immunizations

All mouse experiments were conducted in accordance with procedures approved by the institutional animal care and use committee. Female C57BL/6 or BALB/C mice (purchased from Charles River, Wilmington, MA) were maintained in specific pathogen-free conditions and entered experiments at 6-12 weeks of age unless otherwise indicated. Mice were immunized by intramuscular injection of vaccine delivered in a total volume of 50 μl in the thigh.

#### Pigtail macaque study

Five adult male pigtail macaques were used in these studies (aged 3-6 years, weight 5-13 kg). All animals received a previous Hepatitis B virus (HBV) DNA and protein vaccine regimen, comprised of HBV core and surface antigens and anti-CD180 ^2^, and were re-enrolled in this study in response to the SARS-CoV-2 pandemic. All animals were housed at the Washington National Primate Research Center (WaNPRC), an accredited by the American Association for the Accreditation of Laboratory Animal Care International (AAALAC), and as previously described ^3^. All procedures performed on the animals were with the approval of the University of Washington’s Institutional Animal Care and Use Committee (IACUC).

Blood was collected at baseline (week −2 or −1), and at days 10, 14, 28, and 42 post-prime vaccination (Fig. 5A). Blood was also collected 10 days post-boost (38 days post-prime) in the 50μg vaccinated animals. Serum and plasma were collected and PBMCs were isolated from whole blood as previously described ^4^. Animals were sedated with an intramuscular injection (10 mg/kg) of ketamine (Ketaset^®^; Henry Schein) prior to blood collection or vaccination. Animals were observed daily for general health (activity, appetite) and for evidence of reactogenicity at the vaccine inoculation site (swelling, redness). They also received full physical exams including temperature and weights measurements at each study timepoint. None of the animals became severely ill during the course of the study and none required euthanasia.

#### Pigtail macaque immunization

LION and repRNA-CoV2S were complexed at a nitrogen-to-phosphate molar ratio of 15 in 10mM sodium citrate and 20% sucrose buffer on ice and incubated for at least 30 minutes. The 50μg vaccine was delivered intramuscularly into the quadriceps muscle with one 250 μl injection on weeks 0 and 4. The 250μg vaccine was delivered intramuscularly with five 250μl injections over 4 muscles, 2 in the right quadriceps, 1 in the left quadricep, and 1 each in the left and right deltoids on week 0. All injection sites were shaved prior to injection and monitored post-injection for any signs of local reactogenicity.

#### Serum Chemistries and Complete Blood Counts

Serum chemistries were run on a Beckman Coulter AU 680/5812 system and complete blood counts were determined on a Sysmex XN9000 analyzer by the University of Washington Department of Laboratory Medicine.

#### Antigen-specific antibody responses

Blood was collected from the retro-orbital sinus of immunized mice, or venipuncture of anesthetized macaques, and serum prepared. Antigen-specific IgG, IgG1, IgG2a, and IgG2c responses were detected by enzyme linked immunosorbent assay (ELISA) using a previously described recombinant SARS-CoV-2 S as the capture antigen ^5^. ELISA plates (Nunc, Rochester, NY) were coated with 1 μg/ml antigen or with serial dilutions of purified polyclonal IgG from mouse our monkeys to generate a standard curve in 0.1 M PBS buffer and blocked with 0.2% BSA-PBS. Then, in consecutive order, washes in PBS/Tween, serially diluted serum samples, anti-mouse or-monkey IgG, IgG1, IgG2a, or IgG2c-HRP (Southern Biotech, Birmingham, AL) and TMB then HCL were added to the plates. Plates were analyzed at 405nm (EL_X_808, Bio-Tek Instruments Inc, Winooski, VT). Absorbance values from the linear segment of each serum dilution curve was used to interpolate the standard curve and calculate the IgG concentration present in each sample.

#### SARS-CoV-2 pseudovirus neutralization

Murine leukemia virus (MLV)-based SARS-CoV-2 S-pseudotyped viruses were prepared as previously described ^5, 6^. In brief, HEK293T cells were co-transfected with a SARS-CoV-2 (based on Wuhan-Hu-1 isolate) S-encoding plasmid, an MLV Gag-Pol packaging construct, and the MLV transfer vector encoding a luciferase reporter using the Lipofectamine 2000 transfection reagent (Life Technologies) according to the manufacturer’s instructions. Cells were incubated for 5 hours at 37°C with 8% CO2 with DNA, lipofectamine, and OPTIMEM transfection medium. Following incubation DMEM containing 10% FBS was added for 72 hours. Pseudovirus was then concentrated using a 30kDa Amicon concentrator for 10 minutes at 3,000 x g and frozen at −80C.

BHK cells were plated in 96 well plates for 16-24 hours prior to being transfected with human ACE2 using standard lipofectamine 2000 protocol and incubated for 5 hours at 37°C with 8% CO2 with DNA, lipofectamine, and OPTIMEM transfection medium. Following incubation, DMEM containing 20% FBS was added in equal volume to the OPTIMEM transfection media for 16-24 hours. Concentrated pseudovirus with or without serial dilution of antibodies was incubated for 1 hour at room temperature and then added to the wells after washing 3X with DMEM and removing all media. After 2-3 hours, equal volumes of DMEM containing 20% FBS and 2% PenStrep was added to the cells for 48 hours. Following 48 hours of infection, equal volume of One-Glo-EX (Promega) was added to the cells and incubated in the dark for 5-10 minutes prior to reading on a Varioskan LUX plate reader (ThermoFisher). Measurements were done in duplicate and relative luciferase units (RLU) were recorded.

#### SARS-CoV-2 neutralization

Three-fold (pigtail macaque) or four-fold (human) serial dilutions of heat inactivated serum and 600 plaque-forming units (PFU)/ml solution of SARS-CoV-2/WA/20 (BEI resources) were mixed 1:1 in DPBS (Fisher Scientific) + 0.3% gelatin (Sigma G7041) and incubated for 30 min at 37°C. Serum/virus mixtures were added in duplicate, along with virus only and mock controls, to Vero E6 cells (ATCC) in a 12-well plate and incubated for 1hr at 37°C. Following adsorption, plates were washed once with DPBS and overlayed with a 1:1 mixture of Avicel RC-591 (FMC) + 2 x MEM (ThermoFisher) supplemented with 4% heat-inactivated FBS and Penicillin/Streptomycin (Fisher Scientific). Plates were then incubated for 2 days at 37°C. Following incubation, overlay was removed and plates were washed once with DPBS and then 10% formaldehyde (Sigma-Aldrich) in DPBS was added to cells and incubated for 30 minutes at room temp. Plates were washed again with DPBS and stained with 1% crystal violet (Sigma-Aldrich) in 20% EtOH (Fisher Scientific). Plaques were enumerated and percent neutralization was calculated relative to the virus-only control.

#### Mouse IFN-γ ELISPOT

Spleen and lung lymphocytes were isolated from mice 12 days after the second vaccination. MIAPS4510-Multiscreen plates (Millipore) were coated with rat anti mouse IFN-gamma capture antibody (BD) in PBS and incubated overnight at 4°C. The plates were washed in PBS and then blocked (2h, RT) with RPMI medium (Invitrogen) containing 10% heat inactivated fetal calf serum (Gibco). Lung and spleen cells were plated at 5×10^5^ and 2.5×10^5^ cells/well and stimulated with the SARS-Cov2 S peptide pools (11aa overlapping 15 mer peptides from Genscript) at 1.5 μg/ml/peptide and cultured for 20 hours (37°C, 5% CO2). Biotinylated anti-mouse IFN-gamma antibody (BD) and streptavidin-Alkaline Phosphotase-substrate (Biolegend) were used to detect IFN-gamma secreting cells. Spot forming cells were enumerated using an Immunospot Analyzer from CTL Immunospot profession software (Cellular Technology Ltd).

#### Statistical analyses

Statistical analyses were conducted in Prism (Graphpad) using one-way analysis of variance and Tukey’s multiple comparison test used to compare more than two groups, and either student’s t or Mann Whitney U tests to compare two groups. Statistical significance was considered when the *p*-values were < 0.05.

#### Data availability

Data have been deposited in Figshare: 10.6084/m9.figshare.12385574

### Supplemental Figures

**Supplemental Figure 1.**
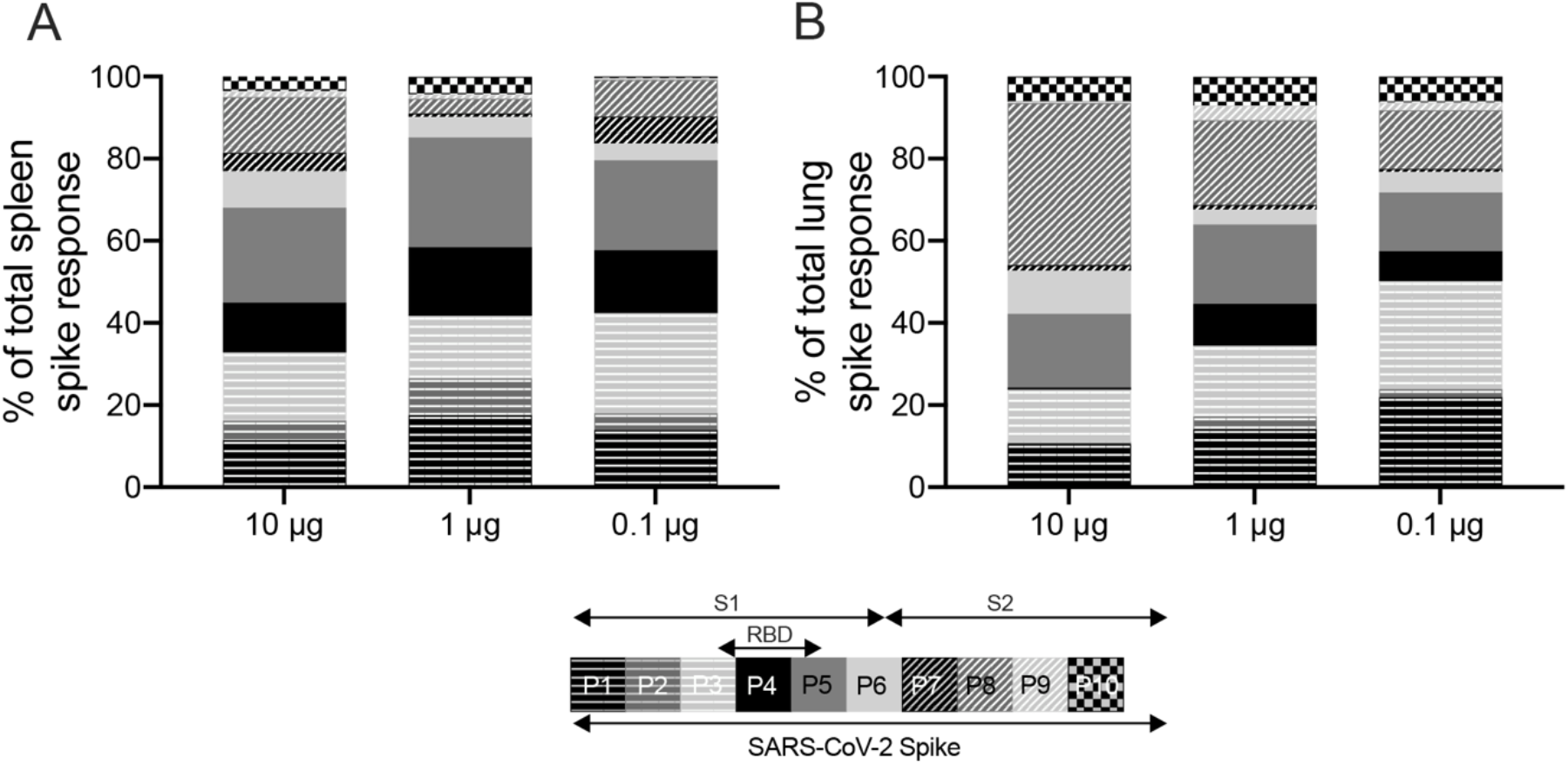
Breadth of T-cell responses in C57BL/6 mice. Six to eight-week old C57BL/6 mice (n=5/group) received 10, 1, or 0.1 μg LION/repRNA-CoV2S via the intramuscular route. On day 28, mice received a booster immunization and 12 days later, (**A**) spleens and (**B**) lungs were harvested and IFN-γ responses were measured by enzyme-linked immune absorbent spot (ELISpot) following stimulation with 10 peptide pools encompassing the entire Spike protein. Each peptide pool consisted of 26-29 15-mer peptides overlapping by 11 amino acids. Data are presented as percent of total spike response.

**Supplemental Figure 2.**
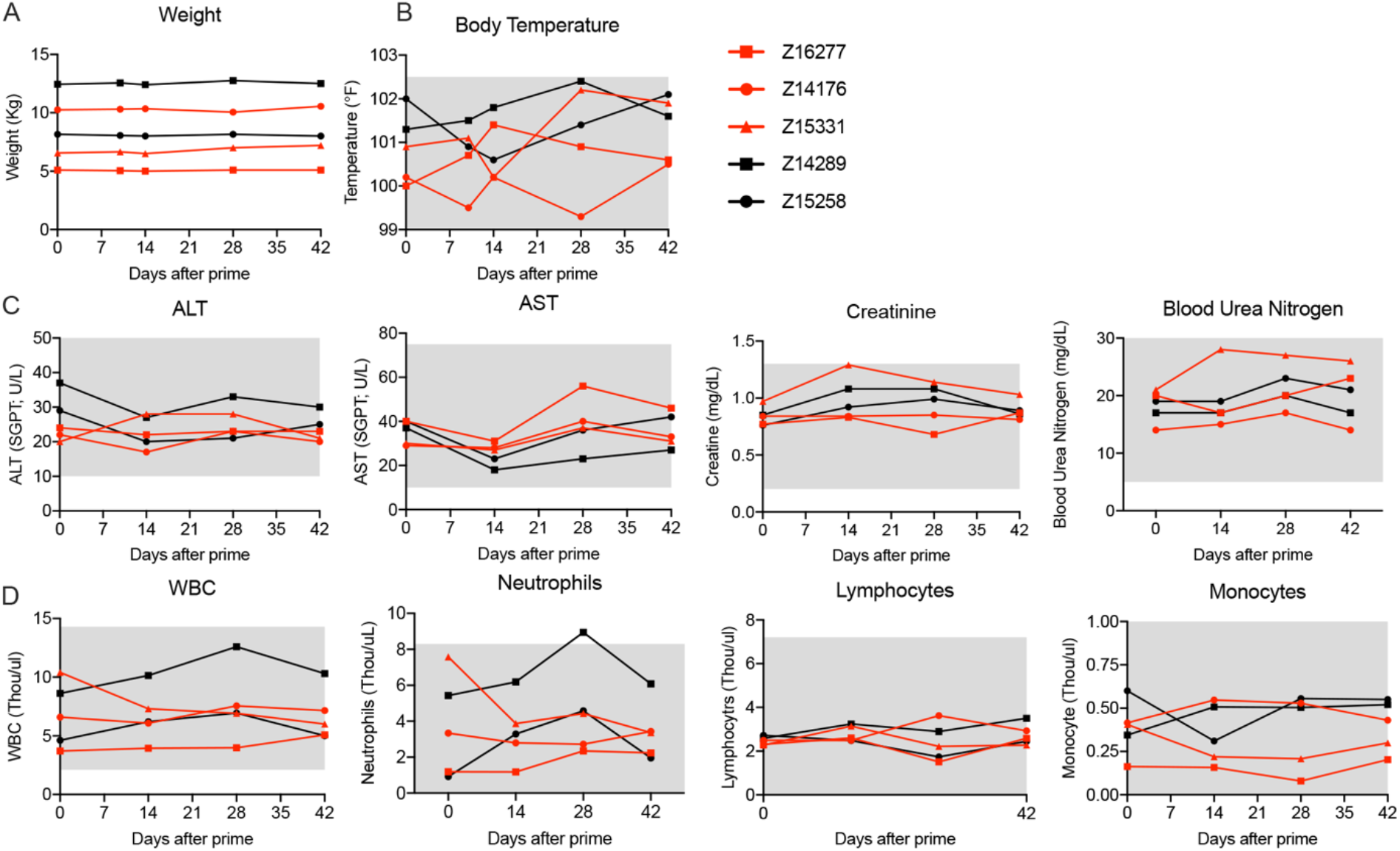
Vaccination did not induce adverse reactions in pigtail macaques. (**A**). Body weight in kg. (**B**) Rectal body temperature in Fahrenheit. (**C**) Serum chemistries. (**D**) Blood complete blood counts (CBC). (**A-D**) Grey shaded areas indicate normal ranges for pigtail macaques.

**Supplemental Figure 3.**
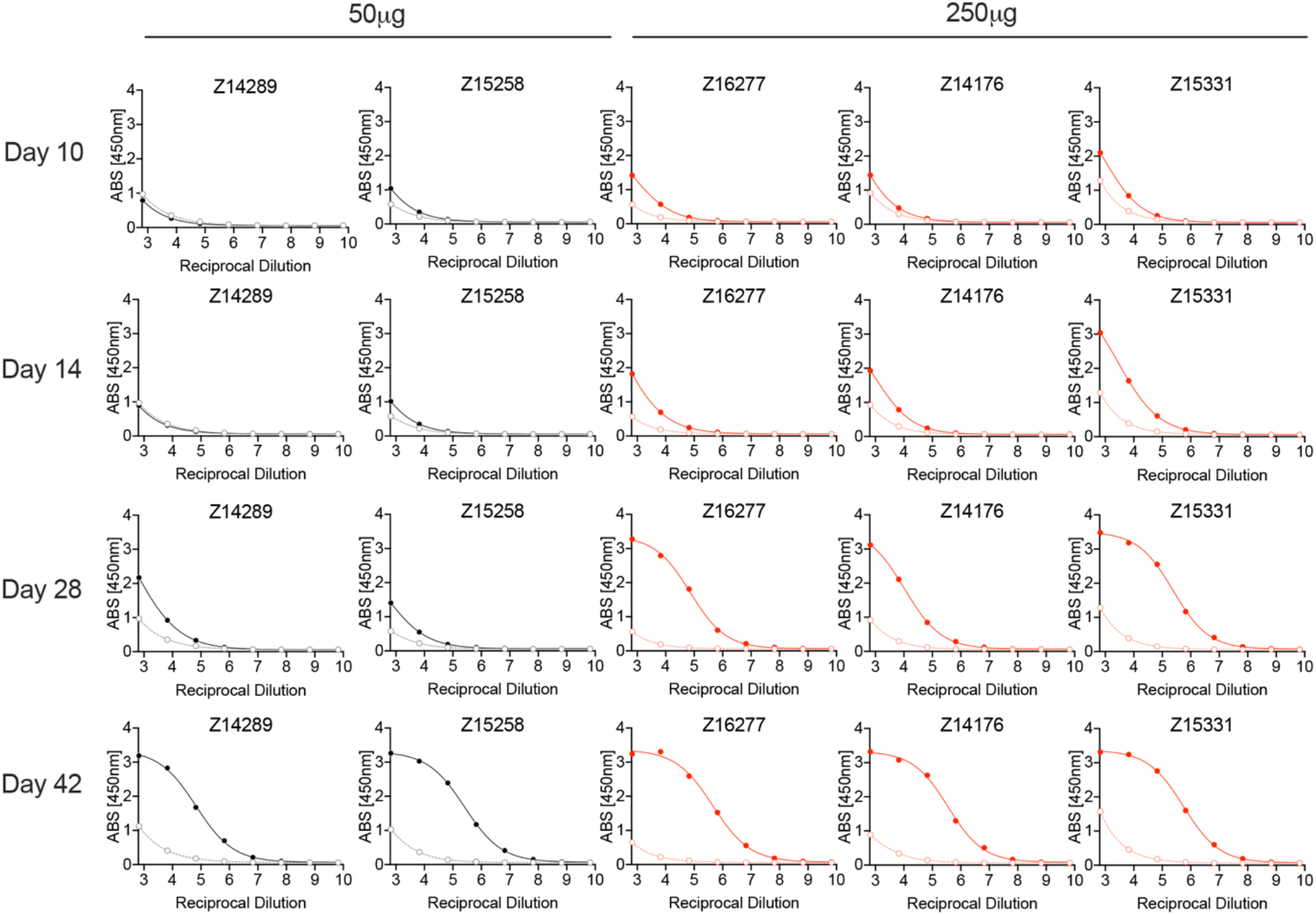
Raw ELISA absorbance values from pigtail macaque study. Recombinant SARS-CoV-2, based on the Wuhan-Hu-1 isolate, was used as the capture antigen and goat anti-monkey IgG-HRP used as the secondary conjugate. Absorbance values were determined at 405nm. Data are presented as pre-immune sera (open circles, dotted line) with post-immune sera (closed circles, solid line). Data are presented as pre-immune sera (open circles, dotted line) with post-immune sera (closed circles, solid line).

**Supplemental Figure 4.**
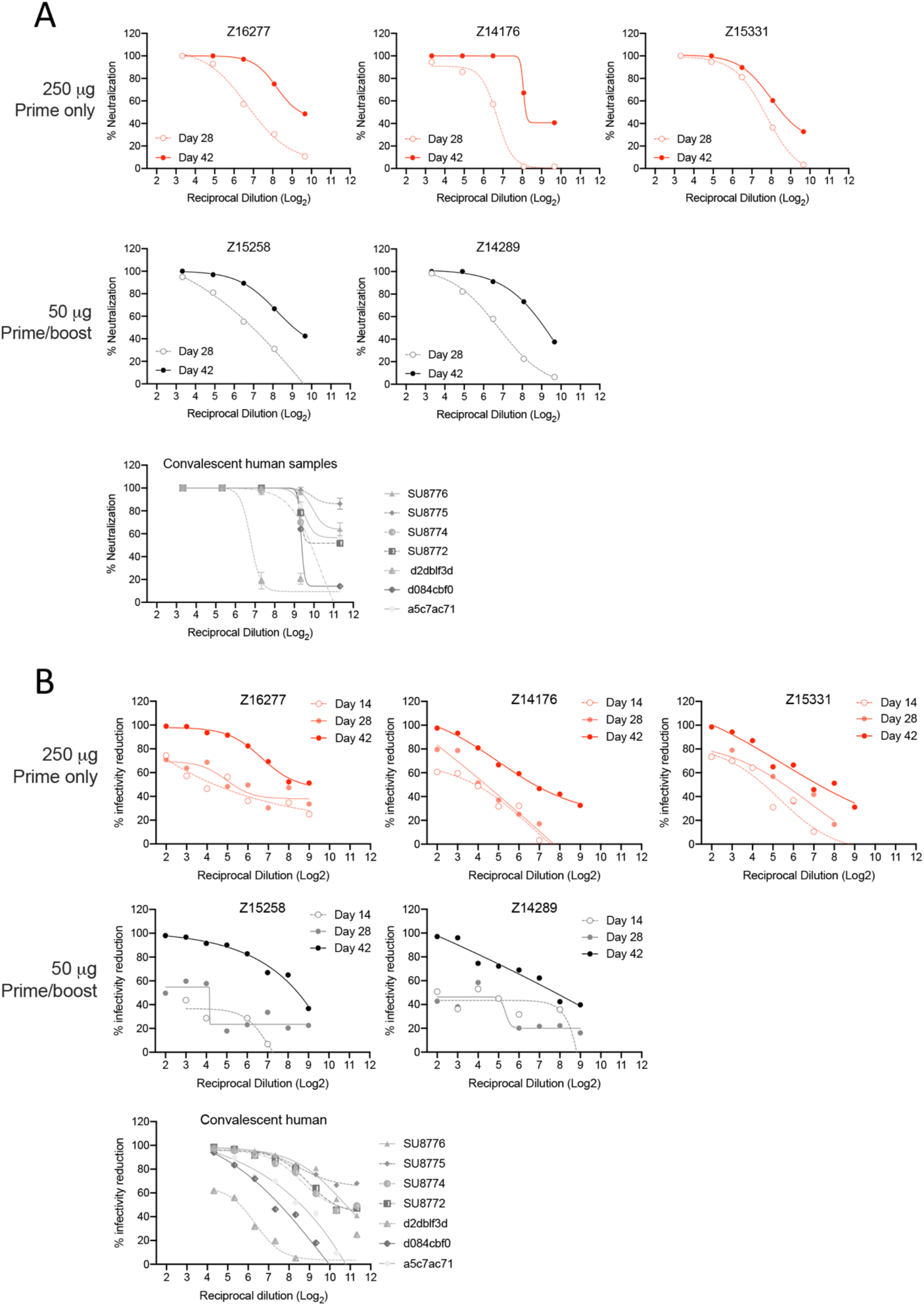
Neutralization curves of pigtail macaque and human samples against (A) SARS-CoV-2/WA/2020 or (B) pseudotyped virus. SARS-CoV-2/WA/2020 neutralization was performed on sera collected from macaques on days 28 and 42 post-primary immunization. Pseudoviral (SARS-CoV-2 Wuhan-Hu-1 pseudotype) neutralization was performed on sera collected from macaques on days 14, 28, and 42 post-primary. (see Sup. Table 1). Both assays were performed alongside sera from 7 convalescent humans collected at various timepoints after their first positive test for SARS-CoV-2 infection.

**Supplemental Table 1.** Convalescent sera from COVID-19 patients

| Sample ID | Days post onset | PRNT<sub>80</sub> |
| --- | --- | --- |
| SU8776 | 20 | 1119 |
| SU8775 | 15 | <20 |
| SU8774 | 21 | 496 |
| SU8772 | unknown | 635 |
| d2db1f3d | 35 | 88 |
| d084cbf0 | 29 | 607 |
| a5c7ac71 | 64 | 658 |

## References

1. Lu, H., Stratton, C. W. & Tang, Y.-W. Outbreak of pneumonia of unknown etiology in Wuhan, China: The mystery and the miracle. J. Med. Virol. 92, 401–402 (2020).

2. Wu, F. et al. A new coronavirus associated with human respiratory disease in China. Nature 579, 265–269 (2020).

3. Wang, C., Horby, P. W., Hayden, F. G. & Gao, G. F. A novel coronavirus outbreak of global health concern. Lancet (London, England) 395, 470–473 (2020).

4. Hamming, I. et al. Tissue distribution of ACE2 protein, the functional receptor for SARS coronavirus. A first step in understanding SARS pathogenesis. J. Pathol. 203, 631–637 (2004).

5. Letko, M., Marzi, A. & Munster, V. Functional assessment of cell entry and receptor usage for SARS-CoV-2 and other lineage B betacoronaviruses. Nat. Microbiol. 5, 562–569 (2020).

6. Walls, A. C. et al. Structure, Function, and Antigenicity of the SARS-CoV-2 Spike Glycoprotein. Cell 181, 281–292.e6 (2020).

7. Walls, A. C. et al. Unexpected Receptor Functional Mimicry Elucidates Activation of Coronavirus Fusion. Cell 176, 1026–1039.e15 (2019).

8. Pinto, D. et al. Structural and functional analysis of a potent sarbecovirus neutralizing antibody. bioRxiv (2020) doi:10.1101/2020.04.07.023903.

9. He, Y., Lu, H., Siddiqui, P., Zhou, Y. & Jiang, S. Receptor-binding domain of severe acute respiratory syndrome coronavirus spike protein contains multiple conformation-dependent epitopes that induce highly potent neutralizing antibodies. J. Immunol. 174, 4908–4915 (2005).

10. Jiang, S., He, Y. & Liu, S. SARS vaccine development. Emerg. Infect. Dis. 11, 1016–1020 (2005).

11. Du, L. et al. The spike protein of SARS-CoV--a target for vaccine and therapeutic development. Nat. Rev. Microbiol. 7, 226–236 (2009).

12. Menachery, V. D. et al. SARS-like WIV1-CoV poised for human emergence. Proc. Natl. Acad. Sci. U. S. A. 113, 3048–3053 (2016).

13. Rockx, B. et al. Structural basis for potent cross-neutralizing human monoclonal antibody protection against lethal human and zoonotic severe acute respiratory syndrome coronavirus challenge. J. Virol. 82, 3220–3235 (2008).

14. Traggiai, E. et al. An efficient method to make human monoclonal antibodies from memory B cells: potent neutralization of SARS coronavirus. Nat. Med. 10, 871–875 (2004).

15. Corti, D. et al. Prophylactic and postexposure efficacy of a potent human monoclonal antibody against MERS coronavirus. Proc. Natl. Acad. Sci. U. S. A. 112, 10473–10478 (2015).

16. Tseng, C.-T. et al. Immunization with SARS coronavirus vaccines leads to pulmonary immunopathology on challenge with the SARS virus. PLoS One 7, e35421 (2012).

17. Honda-Okubo, Y. et al. Severe Acute Respiratory Syndrome-Associated Coronavirus Vaccines Formulated with Delta Inulin Adjuvants Provide Enhanced Protection while Ameliorating Lung Eosinophilic Immunopathology. J. Virol. 89, 2995–3007 (2015).

18. Thanh Le, T. et al. The COVID-19 vaccine development landscape. Nature reviews. Drug discovery (2020) doi:10.1038/d41573-020-00073-5.

19. Shang, W., Yang, Y., Rao, Y. & Rao, X. The outbreak of SARS-CoV-2 pneumonia calls for viral vaccines. npj Vaccines 5, 18 (2020).

20. Xiong, C. et al. Sindbis virus: an efficient, broad host range vector for gene expression in animal cells. Science (80-.). 243, 1188 LP – 1191 (1989).

21. Zhou, X. et al. Self-replicating Semliki Forest virus RNA as recombinant vaccine. Vaccine 12, 1510–1514 (1994).

22. Ljungberg, K. & Liljeström, P. Self-replicating alphavirus RNA vaccines. Expert Rev. Vaccines 18, 1–18 [Epub ahead of print] (2014).

23. Frolov, I. et al. Alphavirus-based expression vectors: strategies and applications. Proc. Natl. Acad. Sci. 93, 11371–11377 (2002).

24. Bredenbeek, P. J., Frolov, I., Rice, C. M. & Schlesinger, S. Sindbis virus expression vectors: packaging of RNA replicons by using defective helper RNAs. J. Virol. 67, 6439–6446 (1993).

25. Liljestrom, P. & Garoff, H. A new generation of animal cell expression vectors based on the Semliki Forest virus replicon. Biotechnology. (N. Y). 9, 1356–1361 (1991).

26. Pushko, P. et al. Replicon-helper systems from attenuated Venezuelan equine encephalitis virus: expression of heterologous genes in vitro and immunization against heterologous pathogens in vivo. Virology 239, 389–401 (1997).

27. Berglund, P., Smerdou, C., Fleeton, M. N., Tubulekas, I. & Liljestrom, P. Enhancing immune responses using suicidal DNA vaccines. Nat. Biotechnol. 16, 562–565 (1998).

28. Jensen, S. & Thomsen, a. R. Sensing of RNA Viruses: a Review of Innate Immune Receptors Involved in Recognizing RNA Virus Invasion. J. Virol. 86, 2900–2910 (2012).

29. Vogel, A. B. et al. Self-Amplifying RNA Vaccines Give Equivalent Protection against Influenza to mRNA Vaccines but at Much Lower Doses. Mol. Ther. 26, 446–455 (2018).

30. Strauss, J. H. & Strauss, E. G. The alphaviruses: gene expression, replication, and evolution. Microbiol. Rev. 58, 491–562 (1994).

31. Kinney, R. M. et al. Attenuation of Venezuelan equine encephalitis virus strain TC-83 is encoded by the 5’-noncoding region and the E2 envelope glycoprotein. J. Virol. 67, 1269–1277 (1993).

32. Atasheva, S. et al. Pseudoinfectious Venezuelan Equine Encephalitis Virus: a New Means of Alphavirus Attenuation. J. Virol. 87, 13 (2016).

33. Erasmus, J. H. et al. A Nanostructured Lipid Carrier for Delivery of a Replicating Viral RNA Provides Single, Low-Dose Protection against Zika. Mol. Ther. 26, 1–16 (2018).

34. Duthie, M. S. et al. Heterologous Immunization With Defined RNA and Subunit Vaccines Enhances T Cell Responses That Protect Against Leishmania donovani. Front. Immunol. 9, 2420 (2018).

35. Calabro, S. et al. The adjuvant effect of MF59 is due to the oil-in-water emulsion formulation, none of the individual components induce a comparable adjuvant effect. Vaccine 31, 3363–3369 (2013).

36. Desbien, A. L. et al. Squalene emulsion potentiates the adjuvant activity of the TLR4 agonist, GLA, via inflammatory caspases, IL-18, and IFN-γ. Eur. J. Immunol. 45, 407–417 (2015).

37. Yu, E. Y. et al. Magnetic Particle Imaging for Highly Sensitive, Quantitative, and Safe in Vivo Gut Bleed Detection in a Murine Model. ACS Nano 11, 12067–12076 (2017).

38. Khandhar, A. P. et al. Evaluation of PEG-coated iron oxide nanoparticles as blood pool tracers for preclinical magnetic particle imaging. Nanoscale 9, 1299–1306 (2017).

39. Bauer, L. M., Situ, S. F., Griswold, M. A. & Samia, A. C. S. High-performance iron oxide nanoparticles for magnetic particle imaging - guided hyperthermia (hMPI). Nanoscale 8, 12162–12169 (2016).

40. Zhao, Y., Zhao, X., Cheng, Y., Guo, X. & Yuan, W. Iron Oxide Nanoparticles-Based Vaccine Delivery for Cancer Treatment. Mol. Pharm. 15, 1791–1799 (2018).

41. Zanganeh, S. et al. Iron oxide nanoparticles inhibit tumour growth by inducing pro-inflammatory macrophage polarization in tumour tissues. Nat. Nanotechnol. 11, 986–994 (2016).

42. Khandhar, A. P. et al. Evaluating size-dependent relaxivity of PEGylated-USPIOs to develop gadolinium-free T1 contrast agents for vascular imaging. J. Biomed. Mater. Res. A 106, 2440–2447 (2018).

43. Zhao, J., Zhao, J. & Perlman, S. T cell responses are required for protection from clinical disease and for virus clearance in severe acute respiratory syndrome coronavirus-infected mice. J. Virol. 84, 9318–9325 (2010).

44. Channappanavar, R., Zhao, J. & Perlman, S. T cell-mediated immune response to respiratory coronaviruses. Immunol. Res. 59, 118–128 (2014).

45. Channappanavar, R., Fett, C., Zhao, J., Meyerholz, D. K. & Perlman, S. Virus-Specific Memory CD8 T Cells Provide Substantial Protection from Lethal Severe Acute Respiratory Syndrome Coronavirus Infection. J. Virol. 88, 11034–11044 (2014).

46. Mills, C. D., Kincaid, K., Alt, J. M., Heilman, M. J. & Hill, A. M. M-1/M-2 Macrophages and the Th1/Th2 Paradigm. J. Immunol. 164, 6166–6173 (2000).

47. Bao, L. et al. Reinfection could not occur in SARS-CoV-2 infected rhesus macaques. bioRxiv 2020.03.13.990226 (2020) doi:10.1101/2020.03.13.990226.

48. Gao, Q. et al. Development of an inactivated vaccine candidate for SARS-CoV-2. Science (80-.). 1932, eabc1932 (2020).

49. Bernstein, D. I. et al. Randomized, double-blind, Phase 1 trial of an alphavirus replicon vaccine for cytomegalovirus in CMV seronegative adult volunteers. Vaccine 28, 484–493 (2009).

50. Hubby, B. et al. Development and preclinical evaluation of an alphavirus replicon vaccine for influenza. Vaccine 25, 8180–8189 (2007).

51. Morse, M. A. et al. Phase I study of alphaviral vector (AVX701) in colorectal cancer patients: comparison of immune responses in stage III and stage IV patients. J. Immunother. Cancer 3, P444–P444 (2015).

52. Reap, E. A. et al. Development and preclinical evaluation of an alphavirus replicon particle vaccine for cytomegalovirus. Vaccine 25, 7441–7449 (2007).

53. Tang, F. et al. Lack of Peripheral Memory B Cell Responses in Recovered Patients with Severe Acute Respiratory Syndrome: A Six-Year Follow-Up Study. J. Immunol. 186, 7264–7268 (2011).

54. Li, C. K. et al. T Cell Responses to Whole SARS Coronavirus in Humans. J. Immunol. 181, 5490–5500 (2008).

## Supplemental References

1. Erasmus, J. H. et al. A Nanostructured Lipid Carrier for Delivery of a Replicating Viral RNA Provides Single, Low-Dose Protection against Zika. Mol. Ther. 26, 2507–2522 (2018).

2. Chaplin, J. W., Chappell, C. P. & Clark, E. A. Targeting antigens to CD180 rapidly induces antigen-specific IgG, affinity maturation, and immunological memory. J. Exp. Med. 210, 2135–2146 (2013).

3. Munson, P. et al. Therapeutic conserved elements (CE) DNA vaccine induces strong T-cell responses against highly conserved viral sequences during simian-human immunodeficiency virus infection. Hum. Vaccin. Immunother. 14, 1820–1831 (2018).

4. O’Connor, M. A. et al. Mucosal T Helper 17 and T Regulatory Cell Homeostasis Correlate with Acute Simian Immunodeficiency Virus Viremia and Responsiveness to Antiretroviral Therapy in Macaques. AIDS Res. Hum. Retroviruses 35, 295–305 (2019).

5. Walls, A. C. et al. Structure, function and antigenicity of the SARS-CoV-2 spike glycoprotein. bioRxiv 2020.02.19.956581 (2020) doi:10.1101/2020.02.19.956581.

6. Millet, J. K. & Whittaker, G. R. Murine Leukemia Virus (MLV)-based Coronavirus Spike-pseudotyped Particle Production and Infection. Bio-protocol 6, e2035 (2016).

